# Nirmatrelvir, an orally active Mpro inhibitor, is a potent inhibitor of SARS-CoV-2 Variants of Concern

**DOI:** 10.1101/2022.01.17.476644

**Authors:** Devendra K. Rai, Irina Yurgelonis, Patricia McMonagle, Hussin A. Rothan, Li Hao, Alexey Gribenko, Elizabeth Titova, Barry Kreiswirth, Kris M. White, Yuao Zhu, Annaliesa S. Anderson, Rhonda D. Cardin

**Author notes:** These authors contributed equally to this work (Devendra K. Rai, Irina Yurgelonis). Corresponding Author mail.

## Abstract

New variants of SARS-CoV-2 with potential for enhanced transmission, replication, and immune evasion capabilities continue to emerge causing reduced vaccine efficacy and/or treatment failure. As of January 2021, the WHO has defined five ‘variants of concern’ (VOC): B.1.1.7 (Alpha, α), B.1.351 (Beta, β), P.1 (Gamma, γ), B.1.617.2 (Delta, δ), and B.1.1.529 (Omicron, o). To provide a therapeutic option for the treatment of COVID-19 and variants, Nirmatrelvir, the antiviral component of PAXLOVID™, an oral outpatient treatment recently authorized for conditional or emergency use treatment of COVID-19, was developed to inhibit SARS-CoV-2 replication. Nirmatrelvir (PF-07321332) is a specific inhibitor of coronavirus main protease (Mpro, also referred to as 3CLpro), with potent antiviral activity against several human coronaviruses, including SARS-CoV-2, SARS-CoV, and MERS (Owen et al, Science 2021. doi: 10.1126/science.abl4784). Here, we evaluated PF-07321332 against the five SARS-CoV-2 VOC (α, β, γ, δ,, o) and two Variants of Interest or VOI, C.37 (λ) and B.1.621 (μ), using qRT-PCR in VeroE6 cells lacking the P-glycoprotein (Pgp) multidrug transporter gene (VeroE6 P-gp knockout cells). Nirmatrelvir potently inhibited USA-WA1/2020 strain, and α, β, γ, λ, δ, μ, and o variants in VeroE6 P-gp knockout cells with mean EC_50_ values 38.0 nM, 41.0 nM, 127.2 nM, 24.9 nM, 21.2 nM, 15.9 nM, 25.7 nM and 16.2 nM, respectively. Sequence analysis of the Mpro encoded by the variants showed ~100% identity of active site amino acid sequences, reflecting the essential role of Mpro during viral replication leading to ability of Nirmatrelvir to exhibit potent activity across all the variants.

## INTRODUCTION

Severe acute respiratory syndrome coronavirus 2 (SARS-CoV-2) was first detected in December 2019 and identified as the causative agent of a novel coronavirus respiratory disease 2019 (COVID-19) [1]. The World Health Organization (WHO) declared COVID-19 a Public Health Emergency of International concern on 20 January 2020 [2] and further characterized the disease outbreak as a pandemic on 11 March 2020 [3].

Similar to other coronaviruses, SARS-CoV-2 encodes an RNA virus 3’→5’ exoribonuclease which reduces its mutation rate, but the genome of SARS-CoV-2 still accumulates mutations over time which could impact viral characteristics, such as transmissibility, virulence or escape from the immune system or therapeutics [4–6]. SARS-CoV-2 variants that spread more readily, cause more severe disease and/or reduce neutralization by virus-specific antibodies are classified as variants of concern (VOCs). As of January 11, 2022, the WHO has designated five VOCs: B.1.1.7 (Alpha, α), B.1.351 (Beta, β), P.1 (Gamma, γ), B.1.617.2 (Delta, δ), and B.1.1.529 (Omicron, o). B.1.1.1.37 (Lambda, λ) and B.1.621 (Mu, μ) have been designated as ‘variants of interest,’ VOI [7]. Some SARS-CoV-2 variants differ in replication kinetics, infectivity, and cytopathicity [8] which could affect the antiviral efficacy of a vaccine or drug against them.

Current determinants for infectivity are mutations in the receptor binding domain (RBD) of the Spike protein of SARS-CoV-2, which mediates virus attachment to the cellular ACE2 receptor. As such, RBD mutations affect the transmission of the variants due to changes in RBD affinity for the receptors and can lead to immune escape from neutralizing antibodies (Nab) induced by vaccination and prior infections [9] as well as reduce efficacy of the Spike targeting monoclonal antibodies that are authorized by the FDA for treatment. Therefore, in addition to vaccines, other treatment modalities such as direct acting antivirals can serve as effective therapeutics to control SAR-CoV-2 infections.

Besides the currently approved antiviral drug Remdesivir, several antivirals are at different stages of development/approval. In particular, two orally available antiviral drug candidates, PAXLOVID (Nirmatrelvir in combination with Ritonavir) and Molnupiravir, received Emergency Use Authorization from the FDA in December of 2021 for treatment of COVID-19 patients [10]. The two drugs have distinctly different mechanisms of action: while Molnupiravir acts as a nucleotide analogue whereas Nirmatrelvir targets the highly conserved Mpro protein of SARS-CoV-2 [11].

Mpro cleaves the viral p1a and p1ab polyproteins at multiple junctions to generate a series of proteins critical for virus replication and transcription, including RdRp, the helicase, and Mpro itself [12]. The essential functional importance in virus replication, together with the absence of closely related homologs in humans, identify Mpro as an attractive antiviral drug target. In this study, we evaluated the currently designated VOCs and two VOIs for susceptibility against Nirmatrelvir to determine *in vitro* efficacy against emerging SARS-CoV-2 variants. In agreement with initial reports [13, 14], Nirmatrelvir exhibits potent activity against Omicron and other evaluated variants and provides critical data supporting the application of Nirmatrelvir in reducing the disease burden from COVID-19.

## MATERIALS AND METHODS

### Virus propagation and characterization

The USA-WA1/2020 strain (Cat. No. NR-52281), α variant (Cat. No. 54000), β variant (Cat. No. 54009), γ variant (Cat. No. 54982), and λ variant (Cat. No. 55664) were purchased from BEI Resources. The δ and μ variants SARS-CoV-2 were obtained from the Hackensack Meridian Health Center for Discovery and Innovation. The Omicron variant B.1.1.529 was obtained from the Icahn Mt. Sinai School of Medicine. All SARS-CoV-2 viruses were propagated by infecting 70-90% confluent Vero E6 TMPRSS2 cells in a T225 cell culture flask in viral growth media (DMEM supplemented with 1 mg/ml geneticin and 10% FBS) at 37°, 5% CO_2_. All viruses were harvested at 48 hours post infection (hpi) at which point ~100% cytopathic effect (CPE) was reached. Cytopathic effect (CPE) was monitored and after reaching ~100% CPE, virus was harvested at 48 hours post infection (hpi). Viral titers were determined as 50% tissue culture infectious dose (TCID_50_) using Reed and Muench method [15]. The lineage of virus and genome sequence was confirmed by NGS analysis.

### Drug preparation

Nirmatrelvir (PF-07321332) and Remdesivir (PF-07304826) were synthesize and stored as previously described [11]. The compounds were dissolved in 100% DMSO to prepare 30 mM stock, which was snap frozen and stored at −80°C until further use.

### Cell culture and EC_50_ assay

The Vero E6 cell line consisting of a knockout in the P glycoprotein (P-gp) multidrug transporter gene (MDR1) was generated by Synthego Corp at the request of Pfizer as a contract service (VeroE6 P-gp knock out (KO) cells, manuscript in preparation). The VeroE6-PgP-ko cells were maintained in a complete growth medium (DMEM supplemented with 1% antibiotic – antimycotic and 10% FBS). Prior to running the assay, Vero E6 Pgp KO cells were grown to 80-100% confluence using the procedure described above. Next, a suspension of the cells was prepared by dissociating the cell monolayer with TrypLE and subsequent dilution in viral growth medium (DMEM supplemented with 1% antibiotic – antimycotic, 2% FBS, and 10 mM HEPES buffer) to 20,000 cells/well. Nirmatrelvir and Remdesivir dilutions were made in virus growth medium containing DMSO at a final concentration of 0.5 %. A 2-fold, 10-point serial dilution was made with Remdesivir starting at 250 nM. Nirmatrelvir was diluted similarly, starting at 1.25 μM. The drug dilutions were then mixed with the cells at 1:1 ratio. The cell + drug mixture was then transferred to BSL-3 and mixed 1:1 (V/V) ratio with SARS-CoV-2 at a MOI of 0.041. After a 48-hour incubation at 37°C, in-plate lysis was performed using 2X lysis buffer. The contents of the plate were transferred to a PCR plate for heat treatment to inactivate SARS-CoV-2 and removed from BSL3 to BSL2 and stored at −80°C until RT-qPCR analysis was completed.

### RT-qPCR

For RT-qPCR, 2 μL of lysate was combined with 18 μL of a master mix consisting of primers, probe, and TaqPath™ 1-Step RT-qPCR Master Mix, CG (Invitrogen, A15299) in an Applied Biosystems™MicroAmp™ Optical 96-Well Reaction Plate followed by plate sealing with MicroAmp™ Optical Adhesive Film (Applied Biosystems, 4360954) and loading into the QuantStudio 7 Pro Thermal Cycler for RT-qPCR analysis. The following thermal cycling conditions for RT-qPCR using QuantStudio Design and Analysis software were used: one cycle of each of 25°C for 2 min, 50°C for 15 min, and 95°C for 2 min, followed by 40 cycles of 95°C for 3 s and 60°C for 30 s. The following oligonucleotides were targeting the SARS-CoV-2 non-structural protein 10 (nsp10) were used: Fwd: 5’-TGACCCTGTGGGTTTTACACTTAA-3’, Rev: 5’-CAGCCATAACCTTTCCACATACC-3’. Probe: 5’-6FAMAACACAGTCTGTACCGTCTMGBNFQ-3’. A standard curve of serially diluted RNA amplicon 5’-GCUAAUGACCCUGUGGGUUUUACACUUAAAAACACAGUCUGUACCGUCUGCGGUAUGUGGAAAGGUUAUGGCUGUAGUU-3’ was run on each plate. After the data were acquired, the percent inhibition was calculated against the virus-infected control cells in the absence of drug using the data analysis method described below.

### Data analysis method

Viral RNA copy number for each well was determined by the QuantStudio Design and Analysis software based on the serially diluted RNA amplicon standard curve run on each RT-qPCR plate. Copy number values were used to calculate percent inhibition of viral replication by Remdesivir or Nirmatrelvir in Excel using the calculation: percent inhibition = 100*((No drug copy number-sample copy no.)/No drug copy number). Percent inhibition versus compound concentration was graphed in GraphPad Prism. EC_50_ and EC_90_ values for both compounds for each virus were calculated in GraphPad Prism using the log(inhibitor) vs. response—Variable slope (four parameters) and [Agonist] vs. response-Find ECanything parameters, respectively. For both calculations, the Hill Slope was set to “must be less than 3.”

### Statistical analysis

For the first set of experiments: The log_10_ of EC_50_ of the variant was analyzed with a one-way ANOVA (with USA-WA1 as the fixed effect) and Dunnett test was used to compare the EC_50_ of the α, β, γ, λ, and δ, variants vs the EC_50_ of the SARS-COV-2 (USA-WA1/2020) strain. For the second and third experiments with the μ and o variants: The log_10_ of the EC_50_ of the variant was tabulated. The EC_50_ data for the USA-WA1/2020 was pooled with the first experiment and the geomean taken across the experiments was compared to the EC_50_ for μ and o variants using a 2-population t-test.

## RESULTS AND DISCUSSION

Vero cells express high levels of MDR1 or P-gp activities, making it necessary to include in previous studies an efflux inhibitor in the cell culture antiviral testing assay to decrease the export of compounds [11, 16]. To eliminate the confounding factors that may be introduced by the inclusion of a co-administered P-gp inhibitor in antiviral activity studies, we created a VeroE6 P-gp knockout cell line (manuscript in preparation).

For antiviral evaluation, the VeroE6 P-gp KO cells were infected at a uniform MOI with each of the SARS-CoV-2 variants tested as described in Materials and Methods. As shown in Table 1, Nirmatrelvir inhibited virus replication in VeroE6 P-gp KO cells with mean EC_50_ values of 38.0 nM, 41.0 nM, 127.2 nM, 24.9 nM, 21.2 nM, 15.9 nM, 25.7 nM, and 16.2 nM in the USA-WA1/2020 SARS-CoV-2 strain and the α, β, γ, λ, δ, μ, and o variants, respectively. In this assay, the EC_50_ for the β variant was 3.3x higher and statistically significant, as compared to the USA-WA1/2020 SARS-CoV-2 strain. The EC_90_ values were 203.4 nM, 212.9 nM, 455.6 nM, 153.4 nM, and 127.2 nM, 26.0 nM, 57.4 nM, for USA-WA1/2020 SARS-CoV-2 strain and the α, β, γ, λ, δ and μ variants, respectively (Table 1). The mean EC_50_ and EC_90_ of Remdesivir (an assay control) were in the range 1.9 - 14.8 nM, and 20.8 - 108.5 nM for the different SARS-CoV-2 variants. Based on the observed EC_50_ values, we conclude that Nirmatrelvir has potent antiviral activity against all current VOCs including Omicron and the two variants of interest, Lambda, and Mu. This result was expected since the Mpro active binding site is highly conserved between the different viruses, although the Mpro contains a K90R mutation in the Alpha and Beta variants, a V296I mutation in the Delta variant and a P132H in the Omicron variant. None of these residues, however, are located in the active site of the enzyme, while K90R and V296I substitutions are conserved and are not expected to induce dramatic structural changes. Therefore, it is not surprising that the observed activity of Nirmatrelvir against these variants was relatively similar. In agreement with our report, two independent groups [14, 17] reported that Nirmatrelvir is active against SARS-CoV-2 VOC, further supporting the efficacy of the drug against the circulating SARS-CoV-2 variants, including Omicron.

**Table 1.** EC_50_ and EC_90_ of Nirmatrelvir Against Major SARS-CoV-2 Variants.

| SARS-CoV-2 | Drug | N | Geo Mean EC<sub>50</sub> (nM)<br>(Range) | N | Geo Mean EC<sub>90</sub> (nM)<br>Range |
| --- | --- | --- | --- | --- | --- |
| **Experiment 1** |  |  |  |  |  |
| USA-WA1 | Remdesivir | 3 | 15.4<br>(10.7 – 21.2) | 3 | 32.3<br>(22.4 – 44.7) |
|  | Nirmatrelvir | 3 | 32.2<br>(15.6 – 90.6) | 3 | 370.9<br>(310.6 – 462.7) |
| $\alpha$ Variant | Remdesivir | 3 | 7.0<br>(5.5 – 9.9) | 3 | 52.2<br>(33.1 – 97.4) |
|  | Nirmatrelvir | 3 | 41.0<sup>a</sup><br>(39.1–45.2) | 3 | 212.9<br>(105.7 – 595.1) |
| $\beta$ Variant | Remdesivir | 3 | 14.8<br>(5.2 – 27.9) | 3 | 108.5<br>(47.7 – 205.7) |
|  | Nirmatrelvir | 4 | 127.2<sup>a</sup><br>(39.8 – 220.4) | 4 | 455.6<br>(344.1 – 682.7) |
| $\gamma$ Variant | Remdesivir | 3 | 3.9<br>(1.6 – 10.5) | 3 | 20.8<br>(10.0 – 33.4) |
|  | Nirmatrelvir | 3 | 24.9<sup>a</sup><br>(15.8 – 33.0) | 3 | 153.4<br>(107.0 – 208.6) |
| $\lambda$ Variant | Remdesivir | 4 | 3.3<br>(1.0 – 5.5) | 4 | 25.0<br>(16.5 – 35.0) |
|  | Nirmatrelvir | 4 | 21.2<sup>a</sup><br>(12.2 – 30.8) | 4 | 127.2<br>(60.2 – 481.5) |
| $\delta$ Variant | Remdesivir | 3 | 1.9<br>(0.9 – 4.5) | 3 | 49.6<br>(25.4 – 69.9) |
|  | Nirmatrelvir | 3 | 15.9<sup>a</sup><br>(8.7 – 37.0) | 2 | 26.0*<br>(18.2 – 33.7) |
| **Experiment 2** |  |  |  |  |  |
| USA-WA1<sup>c</sup> | Remdesivir | 6 | 24.0<br>10.7 – 76.6 | 6 | 72.6<br>22.4 – 452.6 |
|  | Nirmatrelvir | 6 | 36.9<sup>b</sup><br>(15.6 – 90.6) | 6 | 268.6<br>104.6 – 462.7 |
| $\mu$ Variant | Remdesivir | 3 | 5.8<br>3.3 – 8.3 | 3 | 27.8<br>11.4 – 60.4 |
|  | Nirmatrelvir | 3 | 25.7<sup>b</sup><br>(21.9 – 30.2) | 3 | 57.4<br>51.4 – 69.2 |
| **Experiment 3** |  |  |  |  |  |
| USA-WA1<sup>d</sup> | Remdesivir | 9 | 22.5<br>(10.7 – 76.6) | 8 | 63.4<br>(22.4–452.6) |
|  | Nirmatrelvir | 8 | 37.9<sup>e</sup><br>(15.6 – 90.6) | 7 | 203.4<br>(89.2 – 462.7) |
| $\omicron$ Variant | Remdesivir | 3 | 3.2<br>(1.9– 7.3) | | N.A. |
|  | Nirmatrelvir | 3 | 16.2<sup>e</sup><br>(9.4 – 30.5) |  | N.A. |

N=number; \*=mathematical average (N=2)
a. EC<sub>50</sub> values for Nirmatrelvir against variants were evaluated using a one-way ANOVA (with USA-WA1 as the fixed effect) and Dunnett test, the p values for the alpha, beta, gamma, lambda and delta variants were 0.9838, 0.0479, 0.9794, 0.8433 and 0.5213, respectively.
- - b. The $\log_{10}$ of the $EC_{50}$ of the variant was tabulated. $EC_{50}$ data for the USA-WA1/2020 was pooled with the first experiment and geomean taken across experiment 1 and 2 and was compared to the $EC_{50}$ for the Mu variant using a 2-population t-test., the p value was 0.3618. - c. $EC_{50}$ and $EC_{90}$ of USA-WA1 strain combined with experiment 1 ( $N = 3 + 3$ ). - d. $EC_{50}$ and $EC_{90}$ of USA-WA1 strain combined with those from experiment 1, 2 and 3 ( $N = 8$ for nirmatrelvir or 9 for Remdesivir). - e. The $\log_{10}$ of the $EC_{50}$ of the variant (Experiment 3) was tabulated. $EC_{50}$ data for the USA-WA1/2020 was pooled with the first and second experiment and geomean (taken across experiment 1, 2 and 3) was compared to the $EC_{50}$ for the Omicron variant using a 2-population t-test., the p value was 0.0441. - N.A. = Not available

## Author contributions

D.K.R., I.Y., R.A.H., and P.M. conducted experiments. Y.Z., R.D.C., A.S.A. D.K.R, I.Y., R.A.H., and P.M designed and interpreted data results. L.H. and A.G. analyzed sequence and provided structural analysis. D.K.R., Y.Z, A.S.A, and R.D.C. contributed to writing of manuscript. B.K., E.T., and K.W. contributed variants and methods.

## Acknowledgments

We are grateful to the following Pfizer colleagues: Kena Swanson for assistance in acquiring variants, Gretchen Dean for data preparation and discussion, and Nataliya Kushnir for the editorial assistance.

